# Protein Contact Map Denoising Using Generative Adversarial Networks

**DOI:** 10.1101/2020.06.26.174300

**Authors:** Sai Raghavendra Maddhuri Venkata Subramaniya, Genki Terashi, Aashish Jain, Yuki Kagaya, Daisuke Kihara

## Abstract

Protein residue-residue contact prediction from protein sequence information has undergone substantial improvement in the past few years, which has made it a critical driving force for building correct protein tertiary structure models. Improving accuracy of contact predictions has, therefore, become the forefront of protein structure prediction. Here, we show a novel contact map denoising method, ContactGAN, which uses Generative Adversarial Networks (GAN) to refine predicted protein contact maps. ContactGAN was able to make a consistent and significant improvement over predictions made by recent contact prediction methods when tested on two datasets including protein structure modeling targets in CASP13. ContactGAN will be a valuable addition in the structure prediction pipeline to achieve an extra gain in contact prediction accuracy.

## Introduction

Protein structure prediction remains as one of the most important and unsolved problems in biology, more specifically in bioinformatics, biophysics, and structural biology. The tertiary structure of proteins can provide indispensable information for understanding the principle of how the protein structure is built and how it carries out biological functions. Knowledge of the protein structure also facilitates developing drug molecules by enabling structure-based drug design^1,2^ and serves as the basis for artificial protein design^3^. Computational protein structure prediction supplements experimental methods for determining structures. As observed in the community-wide protein structure prediction experiments, the Critical Assessment of techniques in protein Structure Prediction (CASP)^4^, the accuracy of prediction methods has significantly improved in the past few years. The main driver behind this accuracy boost is the improvement of residue-residue contact or distance prediction, which is used effectively to guide the construction of protein structure models^5^.

Residue contacts or distances of a protein are predicted from a multiple sequence alignment (MSA) of the protein^6^. Predicting residue contacts from an MSA of the protein is not a recent novel idea. Rather, it has over 20 years of effort by different research groups toward establishing accurate prediction methods. In principle, evolutionary constraints for maintaining residue-residue contacts in a protein structure leave a trace in the MSA of the protein of interest and its homologous proteins. Earlier works applied relatively simple statistical approaches to detect co-related mutation patterns between two columns (i.e. residue pairs) in a MSA^7-9^. The accuracy of contact prediction was substantially improved a few years ago when the so-called co-evolution approaches, which use statistical inference based on the Potts model^10^, were introduced. The methods in this category include CCMPred^11^, Gremlin^12^, EVFold^13^, plmDCA^10^, FreeContact^14,15^, and MetaPSICOV^15^. Further improvement was observed more recently when deep learning, Convolutional Neural Networks (CNN)^16^ and Residual Networks^17^, were applied to the problem. The methods in this category includes DeepCov^18^, RaptorX-contact^19^, DeepContact^20^, and trRosetta^21^.

Although substantial improvement in contact prediction was observed, contact prediction is still far from perfect. Here, we propose ContactGAN, a novel contact map denoising and refinement method using Generative Adversarial Networks (GAN)^22^. GANs have been widely adopted for high-level generation tasks in computer vision with applications including image-to-image translation^23,24^, image super resolution^25^, and image deblurring^26^. ContactGAN takes a contact map predicted by existing methods, which is considered as a low-resolution or noisy input, and outputs an improved map with a higher accuracy over the original map. ContactGAN was trained with predicted noisy contact maps coupled with corresponding native contact maps, which the networks were guided to generate.

We show that we gain a consistent and substantial precision improvement over predicted maps by CCMPred, DeepCov, and DeepContact. When ContactGAN was applied to the current state-of-the-art contact prediction method, trRosetta, it showed an average precision improvement of 1.14% over L/1 long-contacts on the validation dataset we used. ContactGAN also demonstrated consistent improvement when tested on contact predictions made to the CASP13^5^ protein dataset. It was also demonstrated that combining predicted maps computed by different methods improves the accuracy of generated maps.

### ContactGAN

ContactGAN takes a predicted contact map as input and outputs a refined map. ContactGAN adopts the GAN framework, where two networks, a generative and a discriminative network, are trained with sets of predicted (noisy) and corresponding native (i.e. correct) contact maps so that refined maps can be generated by learning patterns from predicted and native maps. A native contact map we use contains binary values, 1 or 0, where 1 indicates that the Cβ atom distance of the corresponding residue pair is 8 Å or shorter. For training and validation, we prepared a dataset of 6640 non-redundant protein structures (see Methods), for which correct contact maps were generated in addition to predicted maps computed by four existing contact map prediction methods: CCMPred, DeepContact, DeepCov, and trRosetta. These four methods are briefly explained in Methods.

**Figure 1** shows the network structure of ContactGAN. **Figure 1a** illustrates the overall architecture. The generator network, illustrated in **Figure 1b**, is a CNN consisting of a 2D convolution layer with 32 channels and a kernel size of 9, which is followed by 9 or 18 ResNet layers, a 2D convolution layer, and finally by a sigmoid layer. 18 ResNet blocks were used when refining contact maps generated by trRosetta and 9 for all the other methods. Each ResNet block contains 2 convolutional layers with 32 channels and skip connections, dropout (with a dropout probability 0.25), PReLU activations^27^, and Instance Normalization as we used a batch size of 1. In the series of the ResNet blocks, we used dilation filters of 1, 2, and 4 dilations for the first 3 blocks, respectively and repeated this unit of 3 blocks in subsequent layers. The final convolutional layer uses a smaller kernel size of 3 with 32 output channels. The last sigmoid layer outputs a contact probability for each pixel in the 2D contact map. We used a stride of 1 uniformly in all the filters in all the layers.

**Figure 1.**
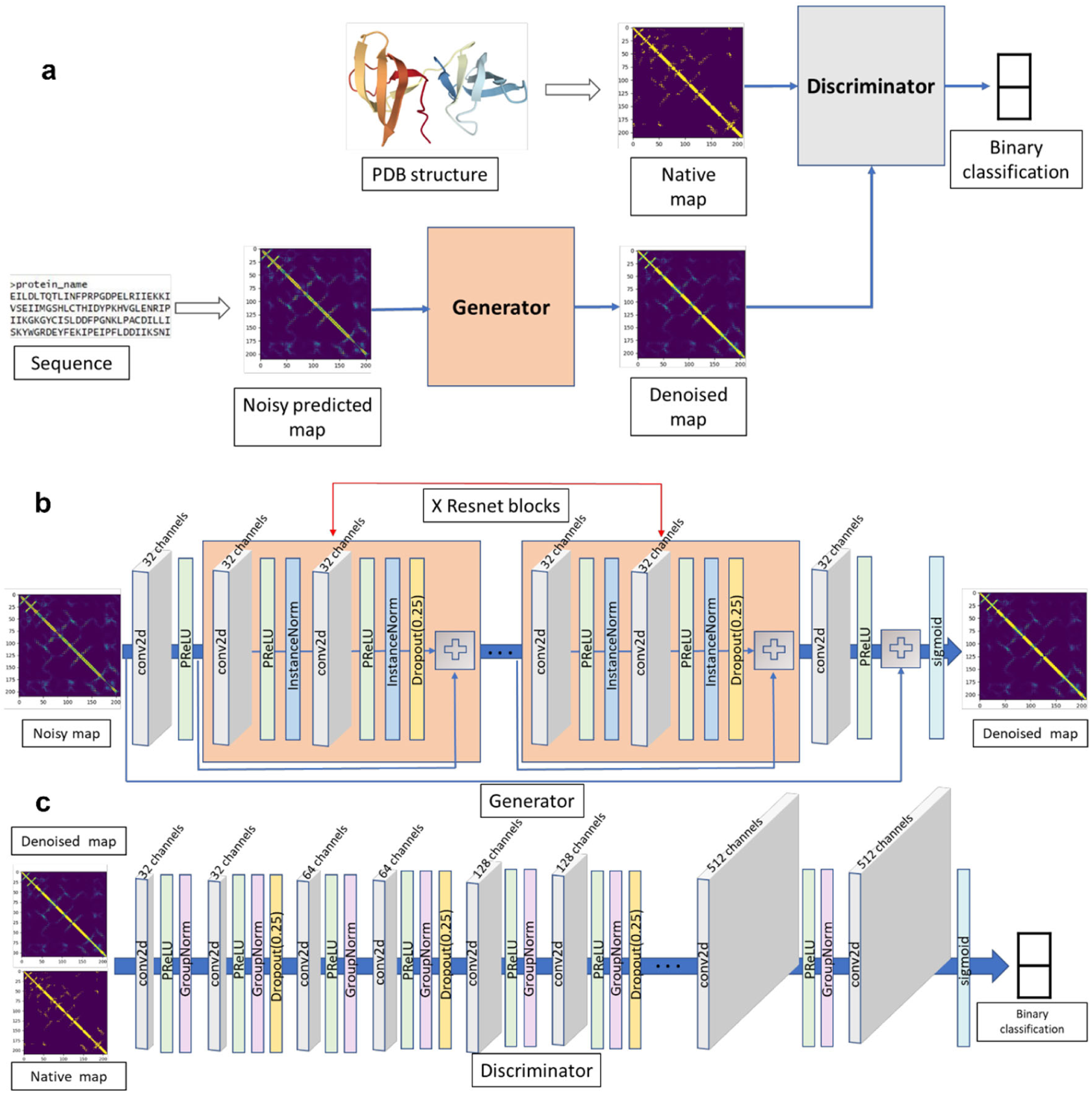
The architecture of ContactGAN. **a**, the overall structure that connects the generator and the discriminator networks. The generator network takes a noisy predicted contact map and outputs a refined map. The discriminator network is to discriminate a generated map by the generator network and the native map, so that the generator is trained to produce indistinguishable maps from native maps by the discriminator. **b**, the detailed network structure of the generator network. X is equal to 18 for handling maps from trRosetta and 9 for the other methods. The blue arrows indicate skip connections, which connect the input of a ResNet block and a plus sign, an operator that simply adds two matrices. **c**, the detailed structure of the discriminator network. See text for more details.

The discriminator network (**Figure 1c)** is a fully convolutional binary classifier. It takes a contact map output from the generator and the corresponding native contact map and classifies the two maps into classes, either native (correct) or predicted. The discriminator network consists of 10 CNNs followed by a softmax layer. The first convolutional layer in the discriminator contains 32 channels, and the count is multiplied by a factor of 2 for each subsequent layer. We used PReLU and group normalization with a group size of 16 in the first convolutional layer and 32 in all other layers.

ContactGAN was trained and validated on the 6,640 proteins in the non-redundant dataset. 6,344 proteins were used for training and 296 were used for validation. ContactGAN was trained separately for each contact prediction method using predicted maps and corresponding native contact maps. Trained models were further tested on 43 protein domains used as targets in CASP13. Note that protein sequence information and other features, such as MSA and secondary structure prediction, were not used in ContactGAN.

### Contact map improvement with ContactGAN

In **Table 1**, we summarize ContactGAN’s performance in improving residue contact map prediction. On the left, results on the validation set in the non-redundant protein dataset are shown while on the right the results on the CASP13 dataset are shown. In this table, we only showed precision considering predicted contacts with the top L/2 and L/1 highest probabilities (L is the length of the protein). Results with more metrics are provided in **Supplementary Table S1, S2, and S3. Supplementary Figure S1** shows improvements of the precision of individual predicted contact maps on the validation and the CASP13 datasets.

**Table 1.** Summary of the improvement of contact map prediction by ContactGAN.

| Method | Validation <sup>a)</sup> |  |  |  | CASP13 |  |  |  |
| --- | --- | --- | --- | --- | --- | --- | --- | --- |
|  | L/2 <sup>b)</sup> |  | L/1 |  | L/2 |  | L/1 |  |
|  | Med+Lg | Lg | Med+Lg | Lg | Med+Lg | Lg | Med+Lg | Lg |
| **CCMpred** | 0.367 | 0.308 | 0.268 | 0.215 | 0.232 | 0.209 | 0.164 | 0.121 |
|  | 0.565 | 0.471 | 0.456 | 0.365 | 0.356 | 0.262 | 0.283 | 0.210 |
| **DeepCov** | 0.636 | 0.521 | 0.503 | 0.391 | 0.394 | 0.287 | 0.32 | 0.231 |
|  | 0.656 | 0.539 | 0.522 | 0.403 | 0.473 | 0.332 | 0.373 | 0.251 |
| **DeepContact** | 0.609 | 0.500 | 0.480 | 0.373 | 0.475 | 0.336 | 0.382 | 0.267 |
|  | 0.649 | 0.540 | 0.521 | 0.409 | 0.516 | 0.372 | 0.410 | 0.283 |
| **C + Dv <sup>c)</sup>** | 0.636 | 0.521 | 0.503 | 0.391 | 0.394 | 0.287 | 0.32 | 0.231 |
|  | 0.683 | 0.577 | 0.557 | 0.443 | 0.500 | 0.363 | 0.396 | 0.283 |
| **C + Dt** | 0.609 | 0.500 | 0.480 | 0.373 | 0.475 | 0.336 | 0.382 | 0.267 |
|  | 0.676 | 0.573 | 0.551 | 0.440 | 0.540 | 0.378 | 0.421 | 0.302 |
| **Dv + Dt** | 0.636 | 0.521 | 0.503 | 0.391 | 0.475 | 0.336 | 0.382 | 0.267 |
|  | 0.691 | 0.579 | 0.558 | 0.441 | 0.527 | 0.39 | 0.420 | 0.309 |
| **C + Dv + Dt** | 0.636 | 0.521 | 0.503 | 0.391 | 0.475 | 0.336 | 0.382 | 0.267 |
|  | 0.706 | 0.600 | 0.576 | 0.463 | 0.554 | 0.397 | 0.443 | 0.310 |
| **trRosetta** | 0.848 | 0.757 | 0.727 | 0.612 | 0.774 | 0.653 | 0.657 | 0.510 |
|  | 0.849 | 0.765 | 0.733 | 0.619 | 0.790 | 0.660 | 0.668 | 0.518 |

**a)** Results of two datasets are shown: On the left, the validation set of 296 proteins in the non-redundant protein dataset; on the right, the CASP13 dataset. **b)** L/2 and L/1 shows precision values when top L/2 or L/1 contact predictions with the highest probabilities were considered. L is the length of the protein. The columns Lg consider only long-range contacts (residue pairs separated by more than 23 residues) while Med+Lg columns consider medium- and long-range contacts (residue pairs separated by more than 11 residues). **c)** Each result show two values: up, original average precision by the existing method; bottom, average precision of denoised contact maps by ContactGAN. In the middle rows, contact maps
of two or three methods were used as input: C, CCMpred; Dv, DeepCov; Dt, DeepContact. When multiple maps were used as input, the highest precision among the existing methods was shown as the original results. For trRosetta, three independent contact predictions were combined, each of which used an MSA with different E-value cutoffs, 0.001, 0.1, and 1.0.

The first three rows show results for individual methods, CCMpred, DeepCov, and DeepContact. ContactGAN made substantial improvements for these methods in all the metrics, which was consistent between the validation set and the CASP13 set. Particularly, the improvements were largest for CCMpred, which had the lowest original precision among the three methods. For CCMpred, ContactGAN improved L/1 Long precision on the validation set from 0.215 to 0.365, an improvement of 69.8%. For the CASP13 set, the improvement for L/1 Long was larger, 73.5% (from 0.121 to 0.210). The improvement ranged from 25.4% to 72.6% for other metrics shown in **Table 1** for CCMpred. The improvement was also consistent for DeepCov and DeepContact, but with smaller improvement margins than observed for CCMpred. For DeepCov, ContactGAN showed an improvement of 3.07% and 8.66% for L/1 Long on the validation and the CASP13 sets, respectively. The corresponding values for DeepContact were 9.65%, and 5.99%, respectively.

The middle rows in **Table 1** present ContactGAN performance using multiple channels, where a pairwise combination of the above three methods and all three methods together were used as input channels. To be able to take two or three contact maps as input, the network architecture of ContactGAN (**Figure 1**) was modified accordingly. The row with “C + Dv” shows the precision values with CCMPred and DeepCov as two input channels for ContactGAN. With these two channels, ContactGAN showed substantial improvement in every evaluation category over the two individual methods. It is interesting to note that the improvement was achieved not only over CCMpred, the method with lower accuracy, but also over the better method, DeepCov. We see similar improvements when CCMPred+DeepContact and DeepCov+DeepContact were used as two-channel inputs. Then, we further extended the use of multi-channels to three channel inputs with CCMpred, DeepCov, and DeepContact altogether (C+Dv+Dt in the table), which resulted in a further improvement over two channel inputs. Improvements by combining additional method(s) are apparent in **Figure 2**, where L/1 Long precision values of each individual method and its combinations with other methods are compared. From originally predicted contact maps predicted by a single method, ContactGAN improved them with a large margin, which was further improved by using two contact maps predicted by different methods (two input channels). Furthermore, an even higher precision was achieved by using three methods as input. Starting from the original prediction by CCMpred, DeepCov, and DeepContact, L/1 Long precisions by the denoised maps using all three methods were improved by 115.3% (from 0.215 to 0.463), 18.4%, and 24.1% on the validation set, and 156.2%, 34.2%, and 16.1% on the CASP13 set, respectively.

**Figure 2.**
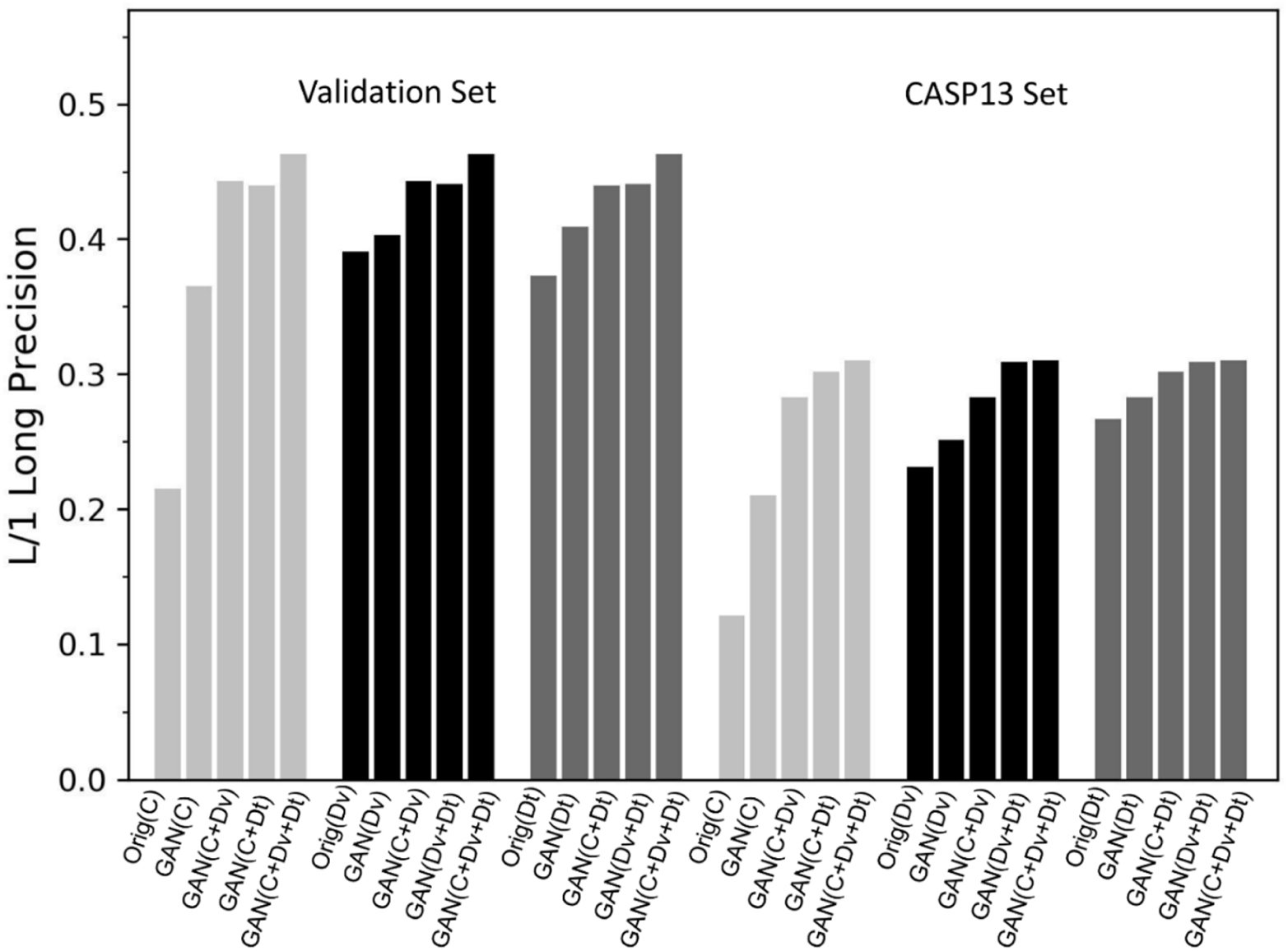
Improvement of L/1 Long precision by using additional predicted contact maps as input channels. Three sets of bars on the left show results for CCMpred (pale gray), DeepCov (black), and DeepContact (gray), respectively, on the validation set. The other three sets on the right are results on the CASP13 set. Orig(C), Orig(Dv), and Orig(Dt) are precision of the original contact maps predicted by CCMpred, DeepCov, and DeepContact, respectively. GAN indicates the precision of denoised maps by ContactGAN. Five bars in a set represent average precision of the original maps, those of the denoised maps by ContactGAN, denoised maps of a combination of two maps, (C+Dv, C+Dt, Dv+Dt), and denoised maps from the combination of three input maps (C+Dv+Dt). Precision values plotted are taken from Table 1.

In the last row of **Table 1** we show the results of the application of ContactGAN to trRosetta, a new method whose performance is one of the best among those published and available^21^. Since the base accuracy of trRosetta is significantly better than the other methods, we combined three different channels of trRosetta, each using different MSAs generated with a sequence E-value cutoff of 0.001, 0.1, and 1.0, respectively, instead of combining with the other methods. Compared with the best prediction among the results with the three E-values, which is 0.001, ContactGAN made consistent improvements for all the metrics. For L/1 Long, ContactGAN improved by 1.14% (from 0.612 to 0.619) and 1.57% (0.510 to 0.518), for the validation set and the CASP13 set, respectively. The performance gains seen on trRosetta are lower than for the other methods, which was as expected since trRosetta provides highly accurate contact maps from the start.

### Contact prediction between secondary structure elements

Next, we investigated which types of contacts were improved by ContactGAN. Particularly, we examined contacts between residues in secondary structure elements, α-helix and α-helix (denoted as α−α below), β-strand and β-strand (β−β), and α-helix and β-strand (α−β). To quantify the change made by ContactGAN, we compared the fraction of correct contacts between secondary structure elements predicted among the top L/1 long-range contacts before and after applying ContactGAN. **Figure 3c** and **3d** show the cases of the three-channel ContactGAN with CCMpred, DeepCov, and DeepContact. For reference, the overall change of L/1 long precision of maps is shown in **Figure 3a** and **3b**. The same plots for the other prediction methods are provided in **Supplementary Figure S2**.

**Figure 3.**
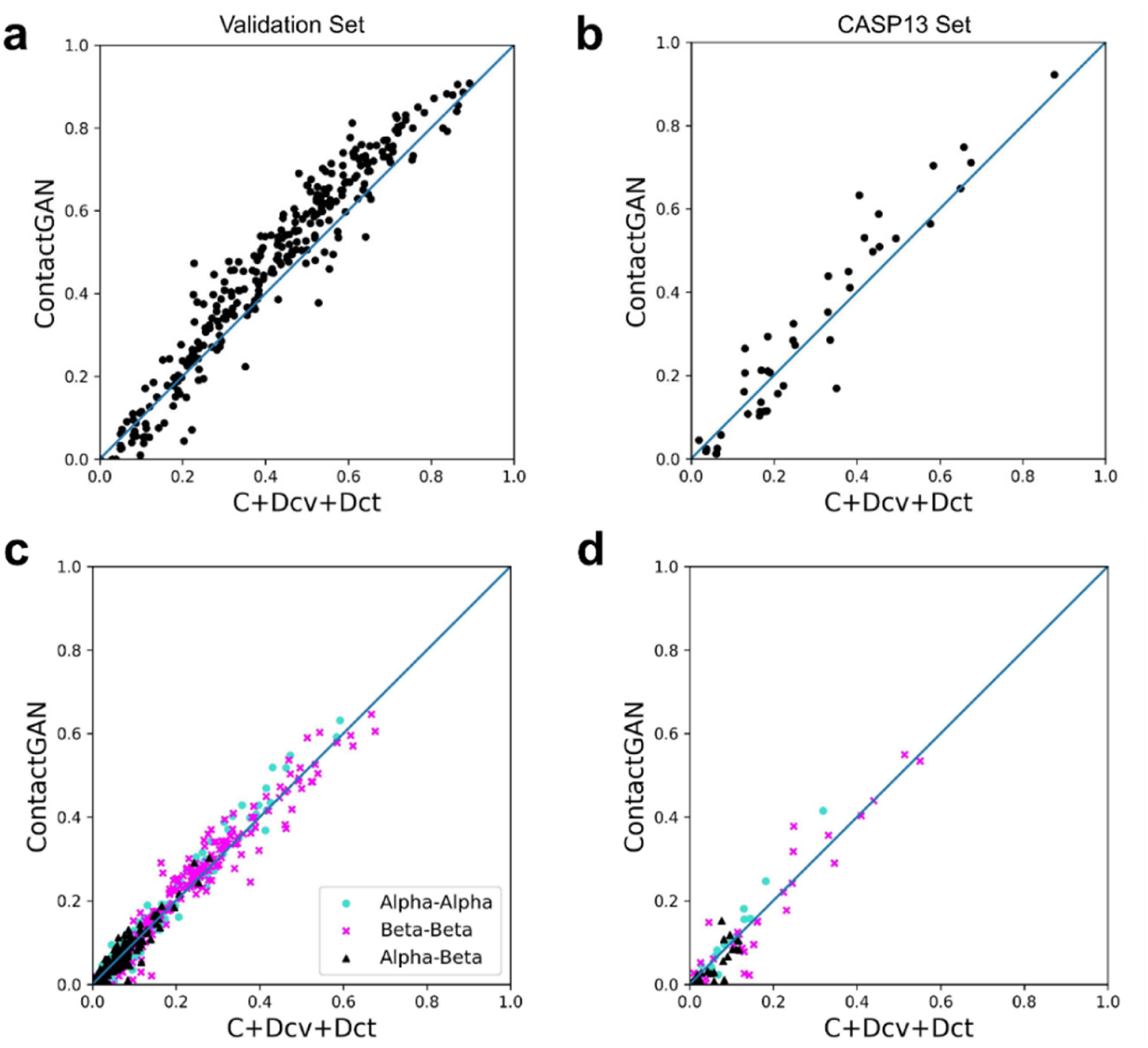
Precision for contacts between secondary structure elements by the three-channel ContactGAN with CCMpred, DeepCov, and DeepContact. **a, b**, the overall L/1 long average precision values before and after applying ContactGAN for each map are plotted. For each map, the x-value shows the highest precision among the three methods, CCMpred, DeepCov, and DeepContact. **a**, the results on the validation set. ContactGAN improved or kept the same precision for 82.1% (243 out of 296) of maps. **b**, the results on the CASP13 set. ContactGAN improved or kept the same precision for 60.5% (26 out of 43) of maps. **c**, The fraction of correct long-range contacts between residues in α-helix and α-helix (α−α; cyan), β-strand and β-strand (β−β; magenta), and α-helix and β-strand (α−β; black) among the top L/1 long predicted contacts are separately plotted. Contacts between residues in loop structure were not considered. The results shown are on the validation set. The fraction of correct contacts between α−α, β−β, and α−β increased or stayed the same for 71.9%, 78.0%, and 72.2% of the maps, respectively. **d**, the results on the CASP13 set. The fraction of correct contacts between α−α, β−β, and α−β increased or stayed the same for 72.1%, 48.8%, and 51.2% of the maps, respectively. Supplemental Figure S2 provides the same type of plots for ContactGAN applied for individual prediction methods.

In **Figure 3c** and **3d**, correct contacts between secondary structure elements share 74.0% and 57.4% among predicted L/1 long contacts in the original prediction for the validation set and the CASP13 set, respectively. On the validation set (**Figure 3c**), all three types of correct secondary structure interactions increased or remained unchanged for about 72 to 78% of the maps. Thus, no preference for the secondary structure interaction type was observed. On the other hand, correct α−α contact predictions were particularly increased in the CASP13 set (**Figure 3d**) compared to the other two types, β−β and α−β contacts, which did not show improvement in about half of the maps. Thus, we can see a tendency that α−α long-range contact predictions have a relatively higher tendency to improve by ContactGAN.

### Examples of improved contact map predictions by ContactGAN

In this section, we show four examples of pairs of contact maps before and after applying ContactGAN. The first example (**Figure 4a**) is a ContactGAN application to a map predicted by CCMpred. For this large protein of 672 amino acids (PDB ID: 3IG5A), the map by CCMpred is covered by noisy predictions with low probability values. The contact probability values of the entire map distributed in a narrow range from 0.0 to 0.17 with the standard deviation of 0.0064 (the diagonal region of the map, i.e. j=i, i+1, i-1 were discarded in these statistics). In contrast, ContactGAN map denoised it into more distinct contact patterns, which yielded a 61.98% improvement in L/1 long-range precision from 0.192 to 0.311. The probability distribution of the denoised map showed minimum and maximum probabilities of 0.0 and 1.0 with a standard deviation of 0.076.

**Figure 4.**
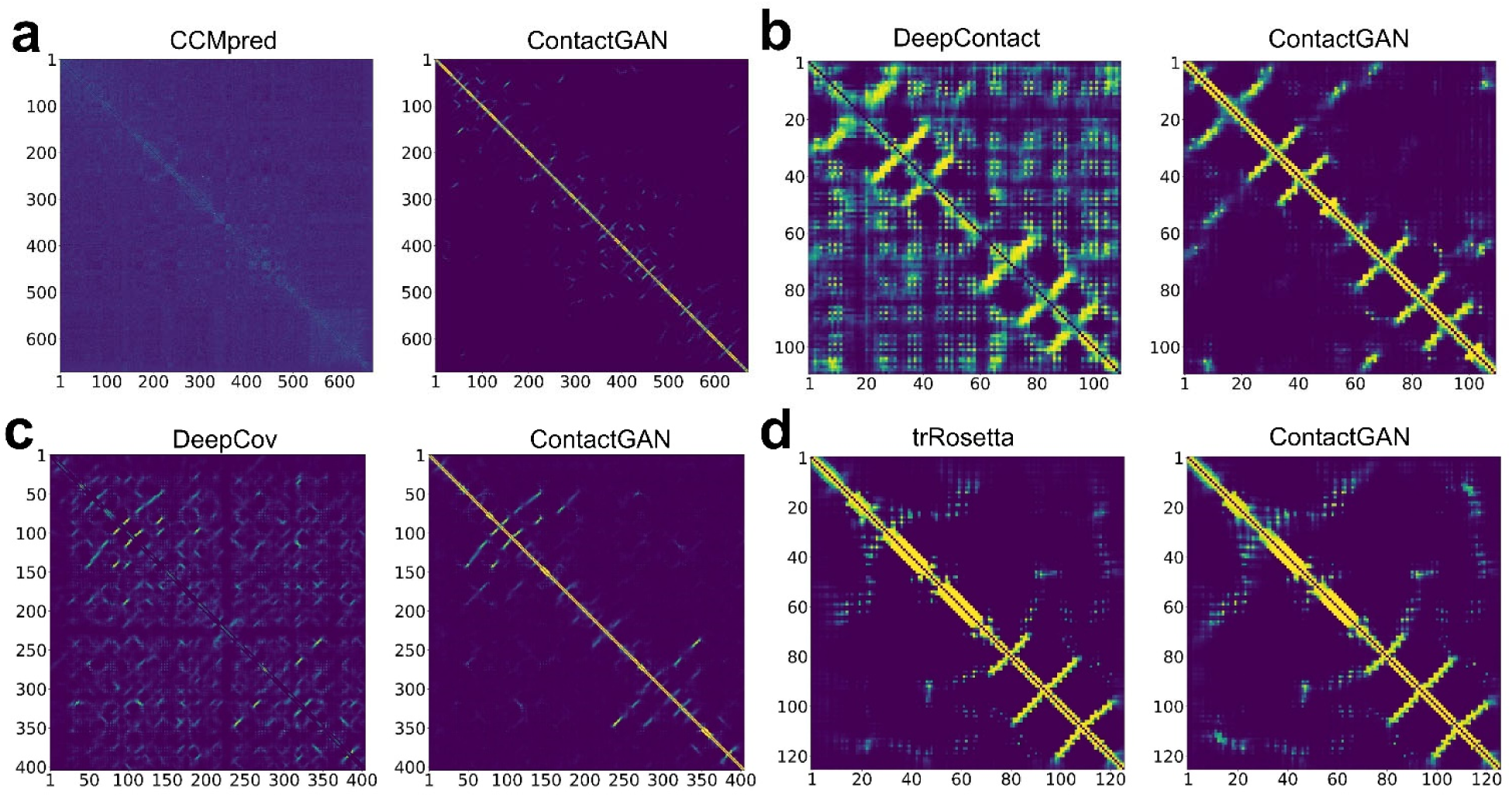
Examples of contact maps before and after applying ContactGAN. For each panel, the map on the left is the original one predicted by an existing method and the map on the right is the denoised map by ContactGAN. The color scale shows predicted probability values of contacts, ranging from dark blue (0.0) to bright yellow (1.0). Contacts with the residue itself along the diagonal line are removed. **a**, a contact map for glutamate-cysteine ligase (PDB ID: 3IG5A; 672 amino acids (aa)) predicted by CCMpred. The L/1 long precision improved from 0.192 to 0.311 by ContactGAN. **b**, a contact map of KOW6-KOW7 domain of human DSIF (PDB ID: 5OHQA, 110 aa) predicted by DeepContact. This map was improved by a two-channel ContactGAN with CCMpred and DeepContact. The comparison here is made with DeepContact because it produced a more accurate map than CCMpred. The L/1 long precision improved from 0.227 to 0.545. **c**, a contact map of a CASP13 target protein, Enterococcal surface protein (CASP ID: T0987, PDB ID:6ORI; 405aa) predicted by DeepCov. It was improved by the three-channel ContactGAN with CCMpred, DeepCov, and DeepContact. This comparison is made with DeepCov because it produced the most accurate map among the three methods for this protein. The L/1 long precision improved from 0.386 to 0.582. **d**, a contact map of a CASP13 target protein, filamentous haemagglutinin family protein (CASP ID: T0968s1, PDB ID:6CP9; 126 aa) by trRosetta using a MSA of E-value 0.001. It was improved by the three-channel ContactGAN with trRosetta with three E-values. The L/1 long precision improved from 0.407 to 0.458.

The next example is the refinement of a predicted contact map by DeepContact for a 110 residue-long β-class protein (**Figure 4b**). On the right panel, we show the output from a two-channel ContactGAN that used predictions from DeepContact and CCMpred as input. Although the original map by DeepContact captures characteristic anti-parallel β-sheet patterns correctly, those are buried in other false-positive contacts with high probability values. ContactGAN was able to clean the strong noise and improved the L/1 long precision from 0.227 to 0.545 over DeepContact, an improvement of 141.1%.

In **Figure 4c**, a map predicted by DeepCov for a 405 residue-long protein in the CASP13 dataset was improved by the three-channel ContactGAN with CCMpred, DeepContact, and DeepCov. Similar to the previous two cases, the original map suffered from high noisy probability values for medium and long-range contacts, which were cleaned by ContactGAN. The L/1 long precision improvement achieved was 50.8% over DeepCov, and 45.1% over DeepContact.

The last example was a contact map by trRosetta (MSA E-value: 10^−3^) refined by the three-channel ContactGAN with trRosetta with three E-values (**Figure 4d**). For this map, a 3-channel input of trRosetta versions was fed to ContactGAN. Compared to the other three cases, the improvements made by ContactGAN visually seem minor; however, they include enhancement of critical very long-range contacts between residues 12-18 and 112-118. These correct contacts were very weakly predicted by trRosetta with the minimum, maximum, and the average values of 0.002, 0.148, and 0.03, respectively, which were strengthened to 0.009, 0.732, and 0.230 for the minimum, maximum, and average probability values, respectively. The precision improvement of L/1 long-range contacts was 12.5% overall.

### Effect of contact map improvement in structure modeling

Improvement in a predicted contact map can often make significant changes in resulting protein structure models that are constructed with the improved map. **Figure 5** shows a few such examples. For the modeling, we used Rosetta^28^ with the constraints from the contact map and also from predicted dihedral angles by SPOT-1D^29^ (see Methods). In the figure, blue and red in a structure model indicate Cα atoms that were modelled within 2.0 Å and over 8.0 Å, respectively, from the correct position measured after superimposing the entire model structure to the native structure.

**Figure 5.**
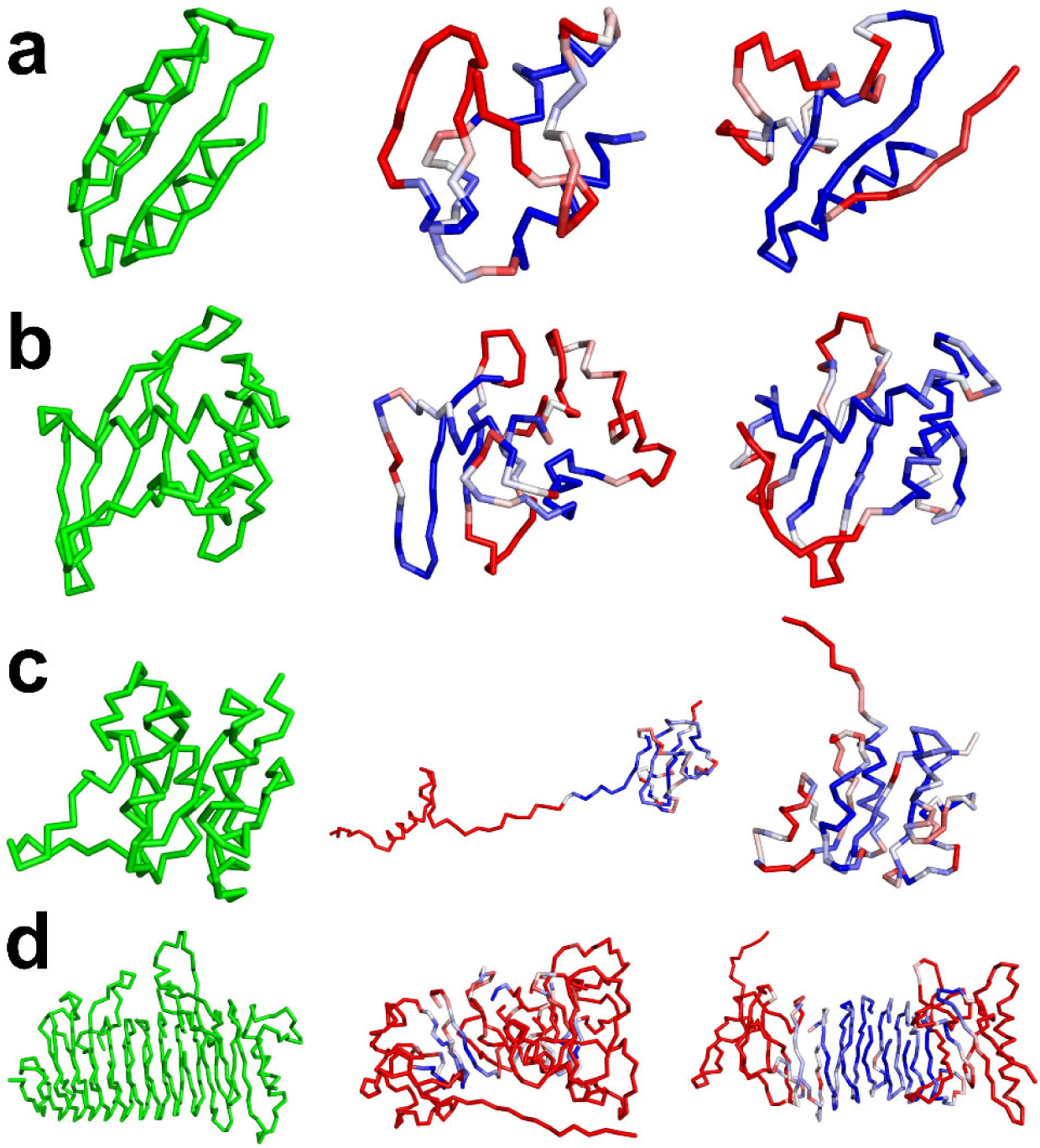
Examples of protein structure models constructed with original and improved contact maps by ContactGAN. Left, the native structure; middle, the protein model constructed with the original contact map; right, the model constructed with the improved map. **a**, an artificially design protein, (PDB ID: 6MSP; 80 amino acids). The original map was predicted by trRosetta. The original contact map had L/1 Long precision was 0.468, which was improved to 0.494 by ContactGAN. With the improved contact map, GDT-TS of the protein model improved from 0.435 (middle) to 0.500 (right). **b**, filamentous haemagglutinin family protein, (PDB ID: 6CP9; 126 aa). The original map was predicted by trRosetta with a L/1 Long precision was 0.407, which was improved to 0.458. With the improved contact map, GDT-TS of the protein model improved from 0.367 (middle) to 0.456 (right). **c**, N-terminal domain of KaiA, (PDB ID: 1M2EA; 135 aa). The original map predicted by trRosetta with a L/1 Long precision of 0.748, which was improved to 0.859. GDT-TS of the protein model improved from 0.322 (middle) to 0.415 (right). **d**, insulin fructotransferase (PDB ID: 2INUA; 410 aa). The original map was predicted by DeepCov with a L/1 Long precision of 0.587, which was improved to 0.652. GDT-TS of the protein model improved from 0.132 (middle) to 0.304 (right).

The first example, **Figure 5a**, is a relatively simple structure with a β-sheet and a layer of two helices. For this protein, ContactGAN improved the L/1 Long precision of the contact map from 0.468 to 0.494. The model using the original map did not have the β-sheet (the middle panel), which was substantially improved by using the denoised map by ContactGAN. The GDT-TS^30,31^ score of the models improved from 0.435 to 0.500 by the improved contact map. In the second example of filamentous haemagglutinin family protein (**Figure 5b**), the improved contact map made possible to correctly fold the core of the structure, which includes a β-sheet and two α-helices. For the protein structure in **Figure 5c**, a drastic effect was caused by the improved contact map, where the over 50-residue long C-terminal tail was now folded. GDT-TS was improved from 0.322 to 0.415 by this change. The last example (**Figure 5d**) is the structure of insulin fructotransferase (PDB ID: 2INUA). The contact map refinement by ContactGAN resulted in substantial improvement of the β-helix structure in the middle of the protein. The GDT-TS increased from 0.132 to 0.304 by the refinement.

## Discussion

In this work, we present ContactGAN, a novel GAN-based denoising framework to push the limit of protein contact prediction. High-quality contact prediction is critical in recent procedures of protein structure prediction. Thus, ContactGAN will be a valuable addition to the structure prediction pipeline to achieve extra gain in the contact prediction accuracy. In principle, ContactGAN can be trained and applied to any contact prediction method. Besides contact prediction, a similar GAN approach will be able to improve other types of structural features used to guide structure modeling, such as main-chain and side-chain angle predictions^21,29^, as well as residue distance predictions^32,33^. Application of GAN to residue distance prediction is left for our future work.

## Acknowledgments

The authors are grateful to Charles Christoffer and Xiao Wang for his help in preparing the manuscript. This work was partly supported by the National Institutes of Health (R01GM123055), the National Science Foundation (DMS1614777, CMMI1825941, MCB1925643, DBI2003635) and the Purdue Institute of Drug Discovery.

## Author Contributions

D.K. conceived and conducted the study. S.R.M.V.S. developed the algorithm and coded ContactGAN. S.R.M.V.S., A.J., and Y.K. made the dataset of predicted contact maps. S.R.M.V.S. carried out the computations. G.T. constructed protein structure models from contact maps. S.R.M.V.S., G.T., and D.K. analyzed the data. S.R.M.V.S. drafted and D.K. edited and finalized the manuscript.

## Methods

### Protein structure and contact map dataset

We prepared a dataset of 6,640 non-redundant protein structures, for each of which a contact map was computed by four existing contact map prediction methods. The predicted contact maps, together with the native (i.e. correct) contact maps, were used for training ContactGAN. A native contact map we use contains binary values, 1 or 0, where 1 indicates that the Cβ atom distance of the corresponding residue pair is 8 Å or shorter.

The protein dataset was constructed based on the PISCES^34^ protein dataset selected with a 25% sequence identity, which was released before May 2018 (i.e. the beginning of CASP13). All these proteins were solved by X-ray or Nuclear Magnetic Resonance (NMR). From the PISCES dataset, proteins longer than 700 amino acid residues or shorter than 25 amino acid residues were discarded. Proteins were also excluded if they contain unknown amino acids in their sequence, have a knot in the structure that was checked by referring to the KnotProt2.0 database^35^, or have consecutive missing residues up to two residues in the structure. Structure gaps up to two residues were filled with the Modeller^36^ automodel protocol.

In addition to this dataset, we used 43 contact prediction (RR) target protein domains in CASP13 as an independent test set. The lengths of the proteins ranged from 80 to 640 (average 243.1). They are available at the CASP13 website (http://predictioncenter.org/casp13/targetlist.cgi).

### Predicting contact maps with four existing methods

We used four existing contact map prediction methods, CCMpred, DeepContact, DeepCov, and trRosetta, to predict contact maps of the proteins in the dataset described above. Input MSAs were generated using DeepMSA^37^. This pipeline collects similar sequences to a target protein sequence in three stages of the database search. We used HHsuite^38^ version 3.2.0 and HMMER^39^ version 3.3 in DeepMSA. For sequence databases, Uniclust30 database^40^ dated October 2017, Uniref90^41^ dated April 2018, and Metaclust_NR database^42^ dated January 2018 were used. These databases were released before CASP13 has started.

CCMpred is a baseline contact prediction method, which uses the Pseudo-Likelihood Maximization (PLM) of direct couplings between pairs of amino acids in a MSA of the target protein^11^. CCMpred has shown more accurate predictions compared to other PLM based methods^11^, such as GREMLIN^12^ and plmDCA^10^. We fed a MSA generated by us using the above-mentioned pipeline.

DeepContact is one of the deep learning-based contact prediction methods^20^. DeepContact uses a CNN, which takes evolutionary coupling information produced by EVFold and CCMpred as well as the secondary structure prediction by PSIPRED^43^ besides an MSA of the target protein. We used the DeepContact code made available at Github by the authors (https://github.com/largelymfs/deepcontact). DeepContact takes a single protein sequence as input and its pipeline generates an MSA and other features for input of CNN and produces a matrix file of predicted residue-residue contacts.

DeepCov is another contact prediction method that uses deep learning^18^. DeepCov uses amino-acid covariance data derived directly from sequence alignments. Unlike DeepContact, DeepCov uses only MSA information as the input of CNNs. We used the code that was available at https://github.com/psipred/DeepCov. Input MSAs were generated by the DeepMSA pipeline described above.

trRosetta uses the ResNet CNN architecture^17^. It was shown that trRosetta had superior performance to other existing methods on the CASP13 dataset. For trRosetta, we generated MSAs at three different E-values in the HHblits search steps of the DeepMSA pipeline, 0.001, 0.1, and 1.0. Since trRosetta outputs predicted distance between residue pairs, we used a distance cutoff of 8Å to decide if two residues are in contact. Since we ran trRosetta with three MSAs of different E-value cutoffs, we obtained three different contact predictions, which were considered as a 3-channel input for ContactGAN.

### Parameter training for ContactGAN

A contact map predicted by a contact map prediction method (e.g. CCMpred), which we hereby refer to as a noisy map, is an input to the generator network of ContactGAN. The output map of the generator network and the corresponding native contact map of the noisy map were then used as inputs to the discriminator network. Out of 6640 pairs of noisy and native contact maps, 6,344 pairs were used for training and 296 were used as validation data.

For training GANs, the generator and discriminator networks are trained together with a min-max game-style objective function given by equation (1):

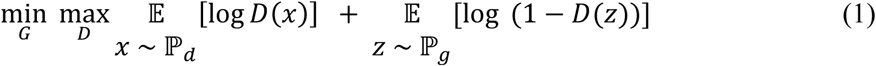

where G and D are parameters of the generator and the discriminator networks of ContactGAN, ℙ_*d*_ is the real (correct) data distribution, ℙ_*g*_ is the generated (fake) data distribution, D(x) and D(z) are predicted probabilities by the discriminator that the real (x) and fake data z are real. The generator receives noisy data as input and generates denoised data. The discriminator then classifies input data to denoised data or real data. The minimax objective ensures that the generator generates good quality denoised data that can fool the discriminator into classifying denoised data as real data.

Following the above common practice for general GAN, the objective function for ContactGAN is formulated as shown in Equation 2, which is a linear combination of a content loss and an adversarial loss:

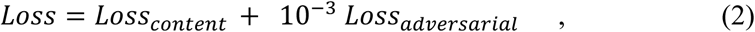

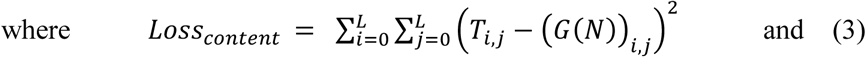

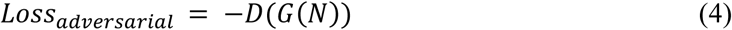

Here, L is the protein sequence length, T corresponds to the native contact matrix (map), and N is the input predicted (noisy) contact matrix, G(N) is the denoised matrix and D(G(N)) is the discriminator’s prediction of the denoised map, which ranges between 0 to 1. We optimize the negative of D(G(N)), as we want to fool the discriminator to consider that the denoised map to be as good as the native map. The content loss is defined by the Mean Squared Error (MSE) between the denoised map and the native map. The adversarial loss is given as the negative softmax probabilities of the discriminator predictions.

We employed the Two Time-scale Update Rule (TTUR)^44^ to use separate learning rates for the generator and the discriminator for stable GAN training. We used learning rates of 0.0001 for the generator and 0.0004 for the discriminator. The batch size was set to 1 as contact maps (i.e. proteins in the dataset) are of different sizes. ContactGAN is trained for 50 epochs. We choose the best performing model on the validation dataset for testing the CASP13 test dataset.

### Building structure models from a contact map

To build protein structure models from the predicted contact map, we used the energy minimization protocol, *MinMover*, in pyRosetta^28^. We added scoring terms that account for Cβ-Cβ contact predictions, which come from an input contact map, and backbone dihedral angle predictions, which was computed by SPOT-1D, in the energy function.

Contact predictions were represented as a flat-harmonic contact potential, *f(x)*, and added to the Rosetta energy:

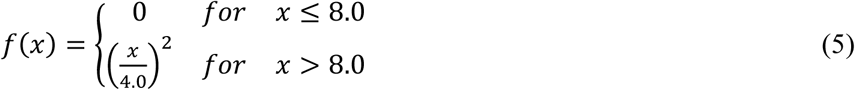

where *x* is the Cβ-Cβ distance between a residue pair. Contact potentials were added to only for the residue pairs that have a contact probability higher than a cutoff value (0.3 and 0.5 were used).

As for the backbone dihedral angle prediction, we added circular harmonic constraints as follows:

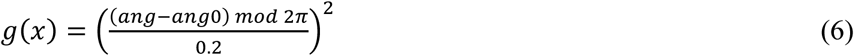

where *ang* is the dihedral angle in the model, *ang0* is a predicted dihedral angle by SPOT-1D. Dihedral angle constraints were added only for residues with a probability of helix or strand that is higher than a cutoff value (0.3 and 0.5 were used).

We used two probability cutoff values, 0.5 and 0.3 for the predicted contact and angle probability cutoff. We used three different folding strategies that differ in the way we consider the short-, medium- and long-range contacts during the energy minimization^21^. Resulting models were relaxed using the *FastRelax* protocol in pyRosetta. For each combination of the probability cutoff and the modeling strategy, we generated 10 models. Thus, in total of 60 models were generated for a target protein. In Figure 5 we reported the model with the largest GDT-TS among the 60 models.

## References

1 Trott, O. & Olson, A. J. AutoDock Vina: improving the speed and accuracy of docking with a new scoring function, efficient optimization, and multithreading. J Comput Chem 31, 455–461 (2010).

2 Shin, W.-H., Christoffer, C. W., Wang, J. & Kihara, D. PL-PatchSurfer2: improved local surface matching-based virtual screening method that is tolerant to target and ligand structure variation. Journal of Chemical Information and Modeling 56, 1676–1691 (2016).

3 Huang, P.-S., Boyken, S. E. & Baker, D. The coming of age of de novo protein design. Nature 537, 320–327 (2016).

4 Kryshtafovych, A., Schwede, T., Topf, M., Fidelis, K. & Moult, J. Critical assessment of methods of protein structure prediction (CASP)—Round XIII. Proteins: Structure, Function, and Bioinformatics 87, 1011–1020 (2019).

5 Shrestha, R. et al. Assessing the accuracy of contact predictions in CASP13. Proteins: Structure, Function, and Bioinformatics 87, 1058–1068 (2019).

6 De Juan, D., Pazos, F. & Valencia, A. Emerging methods in protein co-evolution. Nature Reviews Genetics 14, 249–261 (2013).

7 Ortiz, A. R., Kolinski, A. & Skolnick, J. Fold assembly of small proteins using Monte Carlo simulations driven by restraints derived from multiple sequence alignments. Journal of Molecular Biology 277, 419–448 (1998).

8 Göbel, U., Sander, C., Schneider, R. & Valencia, A. Correlated mutations and residue contacts in proteins. Proteins: Structure, Function, and Bioinformatics 18, 309–317 (1994).

9 Fariselli, P., Olmea, O., Valencia, A. & Casadio, R. Prediction of contact maps with neural networks and correlated mutations. Protein Engineering 14, 835–843 (2001).

10 Ekeberg, M., Lövkvist, C., Lan, Y., Weigt, M. & Aurell, E. Improved contact prediction in proteins: using pseudolikelihoods to infer Potts models. Physical Review E 87, 012707 (2013).

11 Seemayer, S., Gruber, M. & Söding, J. CCMpred—fast and precise prediction of protein residue–residue contacts from correlated mutations. Bioinformatics 30, 3128–3130 (2014).

12 Kamisetty, H., Ovchinnikov, S. & Baker, D. Assessing the utility of coevolution-based residue-residue contact predictions in a sequence- and structure-rich era. Proc Natl Acad Sci U S A 110, 15674–15679 (2013).

13 Marks, D. S. et al. Protein 3D structure computed from evolutionary sequence variation. PloS One 6 (2011).

14 Kaján, L., Hopf, T. A., Kalaš, M., Marks, D. S. & Rost, B. FreeContact: fast and free software for protein contact prediction from residue co-evolution. BMC Bioinformatics 15, 85 (2014).

15 Jones, D. T., Singh, T., Kosciolek, T. & Tetchner, S. MetaPSICOV: combining coevolution methods for accurate prediction of contacts and long range hydrogen bonding in proteins. Bioinformatics 31, 999–1006 (2015).

16 LeCun, Y., Bottou, L., Bengio, Y. & Haffner, P. Gradient-based learning applied to document recognition. Proceedings of the IEEE 86, 2278–2324 (1998).

17 He, K., Zhang, X., Ren, S. & Sun, J. Deep residual learning for image recognition. Proceedings of the IEEE Conference on Computer Vision and Pattern Recognition. 770–778 (2016).

18 Jones, D. T. & Kandathil, S. M. High precision in protein contact prediction using fully convolutional neural networks and minimal sequence features. Bioinformatics 34, 3308–3315 (2018).

19 Wang, S., Sun, S., Li, Z., Zhang, R. & Xu, J. Accurate de novo prediction of protein contact map by ultra-deep learning model. PLoS Computational Biology 13, e1005324 (2017).

20 Liu, Y., Palmedo, P., Ye, Q., Berger, B. & Peng, J. Enhancing evolutionary couplings with deep convolutional neural networks. Cell Systems 6, 65-74. e63 (2018).

21 Yang, J. et al. Improved protein structure prediction using predicted interresidue orientations. Proceedings of the National Academy of Sciences USA (2020).

22 Goodfellow, I. et al. Generative adversarial nets. Advances in Neural Information Processing Systems. 2672–2680 (2014).

23 Isola, P., Zhu, J.-Y., Zhou, T. & Efros, A. A. Image-to-image translation with conditional adversarial networks. Proceedings of the IEEE Conference on Computer Vision and Pattern Recognition. 1125–1134 (2017).

24 Zhu, J.-Y., Park, T., Isola, P. & Efros, A. A. Unpaired image-to-image translation using cycle-consistent adversarial networks. Proceedings of the IEEE International Conference on Computer Vision. 2223–2232 (2017).

25 Ledig, C. et al. Photo-realistic single image super-resolution using a generative adversarial network. Proceedings of the IEEE Conference on Computer Vision and Pattern Recognition. 4681–4690 (2017).

26 Kupyn, O., Budzan, V., Mykhailych, M., Mishkin, D. & Matas, J. Deblurgan: Blind motion deblurring using conditional adversarial networks. Proceedings of the IEEE Conference on Computer Vision and Pattern Recognition. 8183–8192 (2018).

27 He, K., Zhang, X., Ren, S. & Sun, J. Delving deep into rectifiers: Surpassing human-level performance on imagenet classification. Proceedings of the IEEE International Conference on Computer Vision. 1026–1034 (2015).

28 Chaudhury, S., Lyskov, S. & Gray, J. J. PyRosetta: a script-based interface for implementing molecular modeling algorithms using Rosetta. Bioinformatics 26, 689–691 (2010).

29 Hanson, J., Paliwal, K., Litfin, T., Yang, Y. & Zhou, Y. Improving prediction of protein secondary structure, backbone angles, solvent accessibility and contact numbers by using predicted contact maps and an ensemble of recurrent and residual convolutional neural networks. Bioinformatics 35, 2403–2410 (2019).

30 Zemla, A. LGA: a method for finding 3D similarities in protein structures. Nucleic Acids Research 31, 3370–3374 (2003).

31 Zemla, A., Venclovas, Č., Moult, J. & Fidelis, K. Processing and analysis of CASP3 protein structure predictions. Proteins: Structure, Function, and Bioinformatics 37, 22–29 (1999).

32 Xu, J. Distance-based protein folding powered by deep learning. Proceedings of the National Academy of Sciences USA 116, 16856–16865 (2019).

33 Senior, A. W. et al. Protein structure prediction using multiple deep neural networks in the 13th Critical Assessment of Protein Structure Prediction (CASP13). Proteins: Structure, Function, and Bioinformatics 87, 1141–1148 (2019).

34 Wang, G. & Dunbrack, R. L. PISCES: recent improvements to a PDB sequence culling server. Nucleic Acids Research 33, W94–W98 (2005).

35 Dabrowski-Tumanski, P. et al. KnotProt 2.0: a database of proteins with knots and other entangled structures. Nucleic Acids Research 47, D367–D375 (2019).

36 Eswar, N. et al. Comparative protein structure modeling using Modeller. Current Protocols in Bioinformatics 15, 5.6. 1-5.6. 30 (2006).

37 Zhang, C., Zheng, W., Mortuza, S., Li, Y. & Zhang, Y. DeepMSA: constructing deep multiple sequence alignment to improve contact prediction and fold-recognition for distant-homology proteins. Bioinformatics 36, 2105–2112 (2020).

38 Steinegger, M. et al. HH-suite3 for fast remote homology detection and deep protein annotation. BMC Bioinformatics 20, 1–15 (2019).

39 Potter, S. C. et al. HMMER web server: 2018 update. Nucleic Acids Research 46, W200–W204 (2018).

40 Mirdita, M. et al. Uniclust databases of clustered and deeply annotated protein sequences and alignments. Nucleic Acids Research 45, D170–D176 (2017).

41 Suzek, B. E. et al. UniRef clusters: a comprehensive and scalable alternative for improving sequence similarity searches. Bioinformatics 31, 926–932 (2015).

42 Steinegger, M. & Söding, J. Clustering huge protein sequence sets in linear time. Nature Communications 9, 1–8 (2018).

43 Jones, D. T. Protein secondary structure prediction based on position-specific scoring matrices. Journal of Molecular Biology 292, 195–202 (1999).

44 Heusel, M., Ramsauer, H., Unterthiner, T., Nessler, B. & Hochreiter, S. Gans trained by a two time-scale update rule converge to a local nash equilibrium. Advances in Neural Information Processing Systems. 6626–6637 (2017).

