## Supplemental Table S1 for "Protein Contact Map Denoising Using Generative Adversarial Networks"

**Supplementary Table S1A. Summary of improvement by ContactGAN for individual methods on the validation dataset.**

| Method | L/10[a)] | | | L/5 | | | L/2 | | | L | | |
|---|---|---|---|---|---|---|---|---|---|---|---|---|
| | Short | Med | Long | Short | Med | Long | Short | Med | Long | Short | Med | Long |
| **CCMpred[b)]** | 0.366 | 0.419 | 0.481 | 0.274 | 0.322 | 0.418 | 0.171 | 0.202 | 0.308 | 0.122 | 0.138 | 0.215 |
| | 0.569 | 0.600 | 0.618 | 0.479 | 0.522 | 0.582 | 0.321 | 0.368 | 0.471 | 0.205 | 0.245 | 0.365 |
| **Dcv** | 0.733 | 0.705 | 0.725 | 0.618 | 0.605 | 0.657 | 0.406 | 0.425 | 0.521 | 0.250 | 0.280 | 0.391 |
| | 0.741 | 0.715 | 0.735 | 0.625 | 0.619 | 0.671 | 0.410 | 0.437 | 0.539 | 0.252 | 0.286 | 0.403 |
| **Dct** | 0.624 | 0.677 | 0.693 | 0.523 | 0.584 | 0.632 | 0.352 | 0.414 | 0.500 | 0.224 | 0.277 | 0.373 |
| | 0.720 | 0.706 | 0.711 | 0.603 | 0.619 | 0.659 | 0.400 | 0.435 | 0.540 | 0.249 | 0.289 | 0.409 |

Average precision values on the validation dataset of 296 proteins are shown.
**a)** L/k shows precision values when top L/k, k=10, 5, 2, and 1, contact predictions with the highest probabilities were considered. L is the length of the protein. The columns Short consider short-range contacts (residue pairs with a sequence separation of 6-11 residues), Med consider medium-range contacts (residue pairs with a sequence separation of 12-23 residues), and Long consider long-range contacts (residue pairs separated by more than 23 residues). **b)** Each result show two values: up, original average precision by the existing method; bottom, average precision of denoised contact maps by ContactGAN. Dcv, DeepCov; Dct, DeepContact.

**Supplementary Table S1B. Summary of improvement by ContactGAN for individual methods on the CASP13 dataset.**

| Method | L/10 | | L/5 | | L/2 | | L | |
|---|---|---|---|---|---|---|---|---|
| | Med+Lg[a)] | Long | Med+Lg | Long | Med+Lg | Long | Med+Lg | Long |
| **CCMpred** | 0.322 | 0.291 | 0.282 | 0.232 | 0.232 | 0.209 | 0.164 | 0.121 |
| | 0.453 | 0.371 | 0.422 | 0.325 | 0.356 | 0.262 | 0.283 | 0.210 |
| **Dcv** | 0.565 | 0.408 | 0.505 | 0.362 | 0.394 | 0.287 | 0.320 | 0.231 |
| | 0.620 | 0.485 | 0.584 | 0.437 | 0.473 | 0.332 | 0.373 | 0.251 |
| **Dct** | 0.652 | 0.446 | 0.601 | 0.426 | 0.475 | 0.336 | 0.382 | 0.267 |
| | 0.729 | 0.518 | 0.627 | 0.456 | 0.516 | 0.372 | 0.410 | 0.283 |

Average precision values on the CASP13 dataset of 43 protein domains are shown.
**a)** The columns Med+Lg consider medium and long-range contacts (residue pairs separated by more than 11 residues). Other notations are the same as Table S1A.

**Supplementary Table S2A. Summary of improvement by ContactGAN for multi-channel methods on the validation dataset.**

| Method | L/10 | | | L/5 | | | L/2 | | | L | | |
|---|---|---|---|---|---|---|---|---|---|---|---|---|
| | Short | Med | Long | Short | Med | Long | Short | Med | Long | Short | Med | Long |
| **C+Dcv** | 0.733 | 0.705 | 0.725 | 0.618 | 0.605 | 0.657 | 0.406 | 0.425 | 0.521 | 0.250 | 0.280 | 0.391 |
| | 0.751 | 0.738 | 0.746 | 0.633 | 0.647 | 0.694 | 0.419 | 0.457 | 0.577 | 0.255 | 0.298 | 0.443 |
| **C+Dct** | 0.624 | 0.677 | 0.693 | 0.523 | 0.584 | 0.632 | 0.352 | 0.414 | 0.500 | 0.224 | 0.277 | 0.373 |
| | 0.732 | 0.743 | 0.733 | 0.621 | 0.640 | 0.685 | 0.411 | 0.454 | 0.573 | 0.253 | 0.297 | 0.440 |
| **Dcv+Dct** | 0.733 | 0.705 | 0.725 | 0.618 | 0.605 | 0.657 | 0.406 | 0.425 | 0.521 | 0.250 | 0.280 | 0.391 |
| | 0.763 | 0.751 | 0.747 | 0.647 | 0.652 | 0.700 | 0.426 | 0.459 | 0.579 | 0.258 | 0.301 | 0.441 |
| **C+Dcv+Dct** | 0.733 | 0.705 | 0.725 | 0.618 | 0.605 | 0.657 | 0.406 | 0.425 | 0.521 | 0.250 | 0.280 | 0.391 |
| | 0.769 | 0.766 | 0.767 | 0.654 | 0.663 | 0.718 | 0.427 | 0.468 | 0.600 | 0.261 | 0.306 | 0.463 |

The results shown are on the validation dataset. As input, maps from two or three methods were used. To be able to take multiple contact maps, the ContactGAN network architecture was modified to a two-channel or a three-channel GAN. C, CCMpred; Dcv, DeepCov; Dct, DeepContact. For example, C+Dcv indicates that the contact maps from CCMpred and DeepCov were used as input. The highest accuracy among the combined methods was shown as the original results.

**Supplementary Table S2B. Summary of improvement by ContactGAN for multi-channel methods on the CASP13 dataset.**

| Method | L/10 | | L/5 | | L/2 | | L | |
|---|---|---|---|---|---|---|---|---|
| | Med+Lg | Long | Med+Lg | Long | Med+Lg | Long | Med+Lg | Long |
| **C+Dcv** | 0.565 | 0.408 | 0.505 | 0.362 | 0.394 | 0.287 | 0.320 | 0.231 |
| | 0.680 | 0.522 | 0.603 | 0.477 | 0.500 | 0.363 | 0.396 | 0.283 |
| **C+Dct** | 0.652 | 0.446 | 0.601 | 0.426 | 0.475 | 0.336 | 0.382 | 0.267 |
| | 0.726 | 0.562 | 0.647 | 0.495 | 0.54 | 0.378 | 0.421 | 0.302 |
| **Dcv+Dct** | 0.652 | 0.446 | 0.601 | 0.426 | 0.475 | 0.336 | 0.382 | 0.267 |
| | 0.691 | 0.552 | 0.641 | 0.500 | 0.527 | 0.390 | 0.420 | 0.309 |
| **C+Dcv+Dct** | 0.652 | 0.446 | 0.601 | 0.426 | 0.475 | 0.336 | 0.382 | 0.267 |
| | 0.719 | 0.592 | 0.672 | 0.514 | 0.554 | 0.397 | 0.443 | 0.310 |

The results shown are on the CASP13 dataset.

**Supplementary Table S3A. Summary of Improvement by ContactGAN for 3-channel trRosetta on the validation dataset.**

| Method | L/10[a)] | | | L/5 | | | L/2 | | | L | | |
|---|---|---|---|---|---|---|---|---|---|---|---|---|
| | Short | Med | Long | Short | Med | Long | Short | Med | Long | Short | Med | Long |
| **trRosetta (1e-3)[b)]** | 0.868 | 0.876 | 0.889 | 0.758 | 0.794 | 0.855 | 0.490 | 0.569 | 0.757 | 0.283 | 0.358 | 0.612 |
| **trRosetta (1e-1)** | 0.861 | 0.868 | 0.878 | 0.744 | 0.787 | 0.850 | 0.485 | 0.562 | 0.749 | 0.282 | 0.354 | 0.603 |
| **trRosetta (1)** | 0.852 | 0.856 | 0.875 | 0.739 | 0.777 | 0.842 | 0.481 | 0.558 | 0.741 | 0.280 | 0.351 | 0.600 |
| **ContactGAN** | 0.866 | 0.876 | 0.893 | 0.757 | 0.794 | 0.860 | 0.493 | 0.574 | 0.765 | 0.282 | 0.358 | 0.619 |

The results shown are on the validation dataset.

**a)** Top three rows show the average precision by trRosetta which used a MSA with the E-value cutoff specified in the parenthesis. 1e-3: 0.001, 1e-1: 0.1. The last row is the average precision of denoised contact maps by ContactGAN using three trRosetta maps with different E-value cutoffs as input.

**Supplementary Table S3B. Summary of Improvement by ContactGAN for 3-channel trRosetta on the CASP13 dataset.**

| Method | L/10 | | L/5 | | L/2 | | L | |
|---|---|---|---|---|---|---|---|---|
| | Med+Lg | Long | Med+Lg | Long | Med+Lg | Long | Med+Lg | Long |
| **trRosetta (1e-3)** | 0.894 | 0.806 | 0.862 | 0.767 | 0.774 | 0.653 | 0.657 | 0.510 |
| **trRosetta (1e-1)** | 0.919 | 0.816 | 0.879 | 0.751 | 0.780 | 0.637 | 0.648 | 0.500 |
| **trRosetta (1)** | 0.891 | 0.790 | 0.849 | 0.733 | 0.755 | 0.617 | 0.631 | 0.489 |
| **ContactGAN** | 0.909 | 0.829 | 0.878 | 0.770 | 0.790 | 0.660 | 0.668 | 0.518 |

The results shown are on the CASP13 dataset.

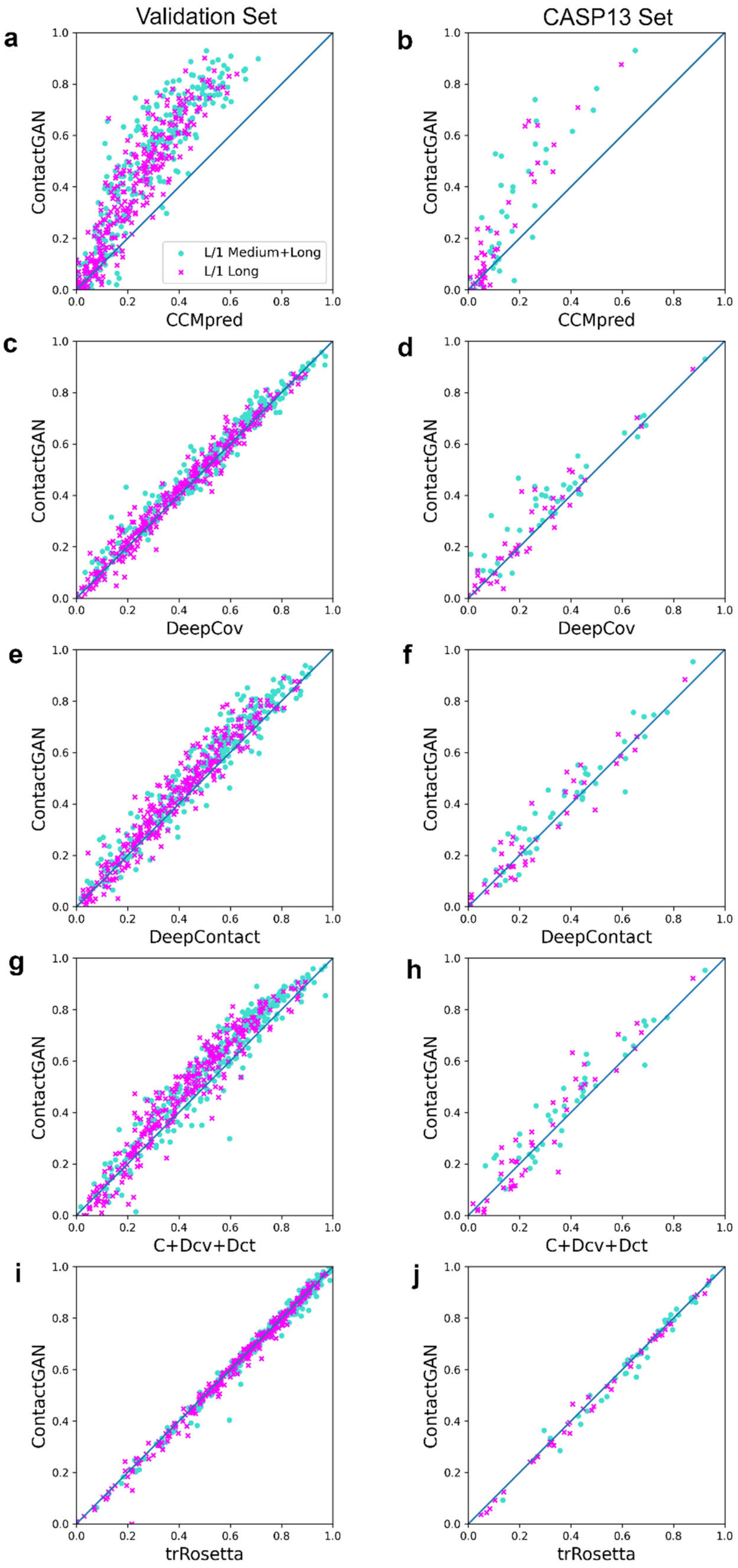
Validation Set
CASP13 Set
a
b
c
d
e
f
g
h
i
j
ContactGAN
CCMpred
DeepCov
DeepContact
C+Dcv+Dct
trRosetta
L/1 Medium+Long
L/1 Long
0.0
0.2
0.4
0.6
0.8
1.0

**Supplementary Figure S1. Precision of ContactGAN relative to the original method in the validation set and the CASP13 set.**

L/1 Long and Medium contacts (circles in cyan) and L/1 Long contacts (crosses in magenta) are plotted for each map in the datasets. **a,** Performance of CCMpred on the validation dataset. **b,** CCMpred on the CASP13 set. **c,** DeepCov on the validation dataset. **d,** DeepCov on the CASP13 set. **e,** DeepContact on the validation dataset. **f,** DeepContact on the CASP13 set. **g,** Performance of the three-channel ContactGAN with CCMPred+DeepContact+DeepCov, on the validation dataset. For the x-axis, for each map the highest precision value among the three methods was plotted. **h,** The three-channel with CCMPred+DeepContact+DeepCov, on the CASP13 dataset. For the x-axis, for each map the highest precision value among the three methods was plotted. **i,** Performance of the three-channel ContactGAN with trRosetta using three E-value cutoffs, 0.001, 0.1, and 1.0 on the validation set. For the x-axis, for each map the highest precision value among the three trRosetta maps was plotted. **j,** the three-channel ContactGAN with trRosetta on the CASP13 dataset. For the x-axis, for each map the highest precision value among the three trRosetta maps was plotted.

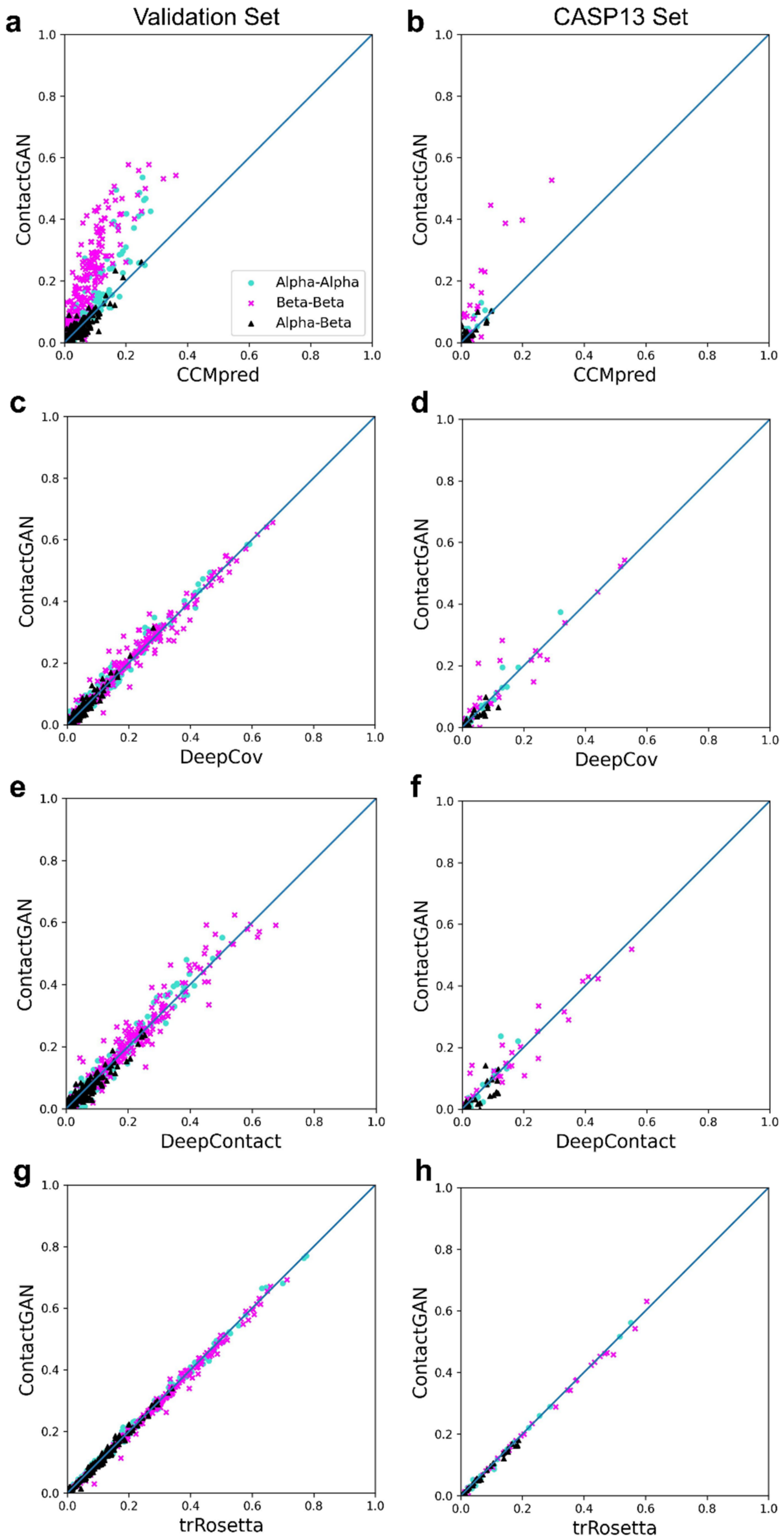


**Supplementary Figure S2. Precision by ContactGAN for contacts between secondary structure elements.**

For each map, the fraction of correct long-range contacts between residues in α-helix and α-helix (α–α; cyan), β-strand and β-strand (β–β; magenta), and α-helix and β-strand (α–β; black) among the top L/1 long predicted contacts are plotted before (x-axis) and after (y-axis) applying ContactGAN. Contacts with a residue in loop structure are not plotted. **a,** the results on predicted maps by CCMpred on the validation dataset. The fraction of correct contacts between α–α, β–β, and α–β increased or stayed the same by ContactGAN for 88.2%, 94.9%, and 73.6% of the maps, respectively. **b,** the results on predicted maps by CCMpred on the CASP13 dataset. Improvement or tie observed for α–α, β–β, and α–β contacts for 93.0%, 86.1%, and 79.1% of the maps, respectively. **c,** the results on predicted maps by DeepCov on the validation dataset. Improvement or tie observed for α–α, β–β, and α–β contacts for 72.9%, 81.7%, and 77.3% of the maps, respectively. **d,** the results on predicted maps by DeepCov on the CASP13 dataset. Improvement or tie observed for α–α, β–β, and α–β contacts for 81.4%, 74.4%, and 69.8% of the maps, respectively. **e,** the results on predicted maps by DeepContact on the validation dataset. Improvement or tie observed for α–α, β–β, and α–β contacts for 72.6%, 74.3%, and 66.9% of the maps, respectively. **f,** the results on predicted maps by DeepContact on the CASP13 dataset. Improvement or tie observed for α–α, β–β, and α–β contacts for 74.4%, 67.4%, and 58.1% of the maps, respectively. **g,** the results on predicted maps by the three-channel trRosetta (three different E-values were used, 0.001, 0.1, and 1.0) on the validation dataset. The x-axis shows the highest accuracy among the three single-channel trRosetta results for each map. Improvement or tie observed for α–α, β–β, and α–β contacts for 81.4%, 64.9%, and 83.8% of the maps, respectively. **h,** the results on predicted maps by the three-channel trRosetta on the CASP13 dataset. Improvement or tie observed for α–α, β–β, and α–β contacts for 83.7%, 65.1%, and 72.1% of the maps, respectively. Figure 3 in the main text shows results for the three-channel ContactGAN with CCMpred, DeepCov, and DeepContact.
